# Dynamic compression schemes for graph coloring

**DOI:** 10.1101/239806

**Authors:** Harun Mustafa, Ingo Schilken, Mikhail Karasikov, Carsten Eickhoff, Gunnar Rätsch, André Kahles

**Author notes:** Equal contribution (Harun Mustafa, Ingo Schilken).

## Abstract

**Motivation:** Technological advancements in high-throughput DNA sequencing have led to an exponential growth of sequencing data being produced and stored as a byproduct of biomedical research. Despite its public availability, a majority of this data remains hard to query to the research community due to a lack of efficient data representation and indexing solutions. One of the available techniques to represent read data is a condensed form as an assembly graph. Such a representation contains all sequence information but does not store contextual information and metadata.

**Results:** We present two new approaches for a compressed representation of a graph coloring: a lossless compression scheme based on a novel application of wavelet tries as well as a highly accurate lossy compression based on a set of Bloom filters. Both strategies retain a coloring with dynamically changing graph topology. We present construction and merge procedures for both methods and evaluate their performance on a wide range of different datasets. By dropping the requirement of a fully lossless compression and using the topological information of the underlying graph, we can reduce memory requirements by up to three orders of magnitude. Representing individual colors as independently stored modules, our approaches are fully dynamic and can be efficiently parallelized. These properties allow for an easy upscaling to the problem sizes common to the biomedical domain.

**Availability:** We provide prototype implementations in C++, summaries of our experiments as well as links to all datasets publicly at https://github.com/ratschlab/graph_annotation.

## 1. Introduction

The revolution of high-throughput DNA sequencing has created an unprecedented need for efficient representations of large amounts of biological sequences. In the next five years alone, the global sequencing capacity is estimated to exceed one exabyte (Stephens *et al*., 2016). While a large fraction of this capacity will be used for clinical and human genome sequencing, such as the 1000 Genomes Project (Auton *et al*., 2015) or the UK10K (Walter *et al*., 2015) effort, that are well suited for reference-based compression methods, the remaining amount is still dauntingly large. This remainder does not only include sequences of model and non-model organisms (Zhang *et al*., 2015) but also community approaches such as whole metagenome sequencing (WMS) (Turnbaugh *et al*., 2007; Ehrlich and Consortium), 2011).

The next logical steps of data integration for genome sequencing projects are assembly graphs that help to gather short sequence reads into genomic contigs and eventually draft genomes. While assembly of a single species genome is already a challenging task (Bradnam *et al*., 2013), assembling a set of genomes from one or many WMS samples is even more difficult. Although preprocessing methods such as taxonomic binning (Dröge and McHardy, 2012) help to reduce its complexity, the task remains a challenge. A commonly used strategy to generate sequence assemblies is based on de Bruijn graphs that collapse redundant sequence information into a node set of unique substrings of length *k* (*k-mers*) and transform the assembly problem into the problem of finding an Eulerian path in the graph (Pevzner *et al*., 2001). Especially in a co-assembly setting, where a mixture of multiple source sequence sets is combined and information in addition to the sequences needs to be stored, *colored* de Bruijn graphs form a suitable data structure, as they allow association of multiple colors with each node or edge (Iqbal *et al*., 2012). A second use case is the application of such graphs for the efficient representation and indexing of multiple genomes, forming a so-called *pan-genome* store (Myers *et al*., 2017).

Owing to the large size, and, subsequently, the excessive memory footprints of such graphs, recent work has suggested compressed representations for de Bruijn graphs based on approximate membership query (AMQ) data structures (Chikhi and Rizk, 2013; Benoit *et al*., 2015) or generalizations of the Burrows-Wheeler transform to graphs (Bowe *et al*., 2012). The recent work on compressed colored de Bruijn graphs has followed this trend. Currently, there exist two distinct paradigms. The first is to compress the complete colored graph in a single data structure while the second proposes two separate (compressed) representations of graph and coloring. Approaches that fall in the first group include *Bloom Filter Tries* (Holley *et al*., 2016) for pan-genome representation, *deBGR* (Pandey *et al*., 2017a) that encodes a weighted de Bruijn graph, or Split Sequence Bloom Trees (Solomon and Kingsford, 2017) that index short read datasets based on a hierarchically structured set of Bloom filters. A very recent addition to this group is *Mantis* (Pandey *et al*., 2017b), that re-purposes the count information in a counting AMQ data structure to represent a set of colors. The second group contains approaches such as *VARI* (Muggli *et al*., 2017), that uses succinct Raman-Raman-Rao or Elias-Fano compression on the annotation vector, and *Rainbowfish* (Almodaresi *et al*., 2017), that additionally takes into account the distribution of the annotations in the graph to achieve better compression rates.

Our contribution falls into the second group and allows for efficient addition and removal as well as editing of individual annotation tracks on an existing graph structure. We present a succinct annotation data structure based on wavelet tries that takes advantage of correlations between columns of the annotation matrix and shows excellent compression rates on a wide range of input data. Moreover, the proposed data structures can efficiently handle dynamic settings where annotation or underlying graph structure are subject to change. For genomics applications, such as encoding of a pan-genome index for read labeling, where a fully exact reconstruction of the annotation is not necessary but an approximate recovery with high accuracy would be sufficient, we also present a probabilistic compression scheme for an arbitrary number of colors. Based on Bloom filters (Bloom, 1970), a data structure for efficient AMQ with a one-sided error, we encode colors as bit vectors and store them in a set of filters. We further reduce the necessary storage requirements of the individual filters by maintaining weak requirements on their respective false-positive rates, which is subsequently corrected for by using neighborhood information in the graph.

After providing a short introduction to our setup and notation in Section 2.1, we give a summary of the graph structure used in our experiments in Section 2.2. We then introduce a lossless color compression scheme based on wavelet tries in Section 2.3.1 and provide a description of a probabilistic encoding that drastically improves the compression rate at a moderate loss of accuracy in Section 2.3.2. Next, in Section 3 we evaluate our proposed strategies in the context of commonly used color encodings on a wide range of different datasets. Finally, we use Section 4 to discuss limitations and give an outlook on future directions.

## 2 Approach

The proposed techniques for color compression take advantage of the underlying sequence graph. Although we impose no restrictions on graph topology, we assume that all nodes in a *linear path* (a directed path in which all nodes have in-degree and out-degree 1) share an identical *coloring* (a set of colors). In this work, we will focus on compressing colorings of de Bruijn graphs constructed on pangenomic and metagenomic datasets.

We implement our reference metagenome as a *colored de Bruijn graph* (cDBG), which consists of a de Bruijn graph constructed from a collection of input sequences (forward and reverse complement) and an *annotation* associated with the *k*-mers generated from these input sequences. We represent this annotation as a binary matrix, where each row corresponds to an edge and each column corresponds to a predefined annotation class. Set bits in this matrix indicate associations of edges with annotation classes.

### 2.1 Preliminaries and notation

Let Σ be an alphabet of fixed size (in the case of genome graphs, Σ = {$, *A, C, G, T, N* }, where $ represents the string terminus). Given a string *s* ∈ Σ*, we use *s*[*i* : *j*] to denote the substring of *s* from index *i* up to and including index *j*, with *i, j* ≥ 1.

Given a bit vector *b* ∈ {0, 1}^*m*^ of length *m*, we use the notation |*b*| to refer to its length, *b*[*i*] to refer to its *i*^th^ character, 1 ≤ *i* ≤ |*b*|, *b*[*j* : *k*] to refer to the bit vector *b*[*j*] … *b*[*k*], *b*[: *k*] to refer to its prefix *b*[1 : *k*], and *b*[*j* :] to refer to its suffix *b*[*j*] … *b*[|*b*|]. The empty vector is denoted *ε*. Finally, given bit vectors *a, b* ∈ {0, 1}^*m*^, we use the notation *a* ⋁ *b* and *a* ⋀ *b* to denote the bitwise OR and AND operators, respectively.

The function rank_0_(*b, j*) counts the occurrences of the character 0 in the prefix *b*[: *j*], while select_0_(*b, j*) returns the index of the *j*^th^ 0 in *b*. The functions rank_1_ and select_1_ are defined analogously for the 1 character. We will use the notation 2^*A*^ to denote the power set of a set *A* and abuse the notation | · | to also denote set cardinalities.

### 2.2 Graph representation

Given an ordering of the edges *E* = (*e*_1_, …, *e*_*n*_) of an underlying graph *G* = (*V, E*) and a set of colors 1, …, *m*, we define the *annotation matrix* 𝒜 ∈ {0, 1}^*n*×*m*^ such that

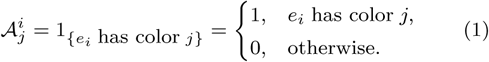

As a proof of concept for the graph coloring presented in this work, and without loss of generality, we use a simple representation of a de Bruijn graph with its edges (the *k*-mers) stored in a hash table.

During construction of the graph, colors are computed for each edge based on the metadata of the input sequences. We assign each unique metadata string to an annotation category with a corresponding positive integer index (*color*). During *k*-mer enumeration, each *k*-mer is assigned to a set of colors encoding its respective metadata categories. We then represent this coloring through a binary vector (*bit vector*) with bits set for the corresponding edge colors. When duplicate *k*-mers are collected to construct graph edges, we combine the *k*-mers’ respective bit vectors via bit-wise OR operations and assign the aggregated coloring to the resulting edge. Alongside the de Bruijn graph, this process results in the encoding of a *graph coloring* as an annotation matrix 𝒜 with *n* rows corresponding to the edges of the graph and *m* columns corresponding to the total number of annotation classes observed during construction. The resulting graph-annotation pair (𝒢, *𝒜*) is a *colored de Bruijn graph*. When the graph is queried, patterns are mapped to a path (a sequence of edges) and, hence, a corresponding sequence of annotation matrix rows.

### 2.3 Graph topology-aided color compression

#### 2.3.1 Loss-less row compression with wavelet tries

For lossless compression of annotation matrices, we propose a novel application of the *wavelet trie* data structure (Grossi and Ottaviano, 2012). Wavelet tries compress tuples of dynamic bit vectors by finding common *segments* (contiguous subsequences) among the encodings of its characters. Briefly, a wavelet trie builds on the concept of a wavelet tree and takes the shape of a compact prefix tree (a binary radix trie, cf. Figure 1 and Suppl. Figure S-1).

**Fig. 1:**
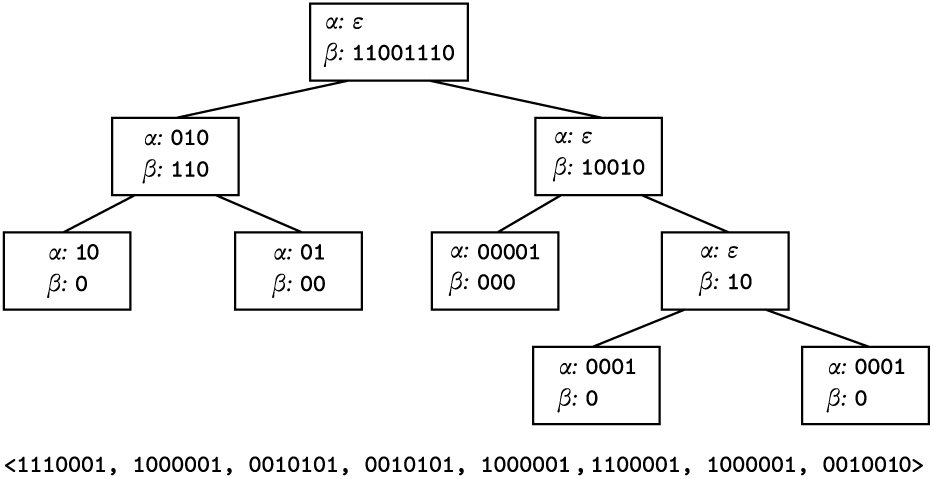
A wavelet trie constructed for a tuple of bit vectors. At a node, the common prefix of the bit vectors is extracted and the next significant bit is used to assign the bit vector suffices to that node’s children. A node becomes a leaf when all bit vectors assigned to it are equal. Index queries are resolved by traversing the tree and performing rank operations on the assignment bit vectors.

In the context of genome graph coloring, we employ wavelet tries to compress the rows of the annotation matrix to allow for dynamic updates in its rows and columns. We employ a construction strategy based on wavelet trie merging (Grossi and Ottaviano, 2012; Böttcher *et al*., 2017), but in a parallel fashion. Since wavelet tries were originally conceived to compress binary encodings of strings (where the null terminal character has an encoding), the assumption that the end of a sequence is marked by a specific subsequence no longer holds in our application. Thus, our construction and merging algorithms are adapted to take this fact into account.

##### Construction

The wavelet trie encoding the annotation matrix 𝒜 ∈ {0, 1} ^*n*×*m*^ is constructed recursively and is a binary tree (Figure 1) with nodes *V*_*T*_ of the form

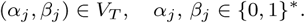

The *α*_*j*_ are referred to as the *longest common prefices* (LCPs) and the *β*_*j*_ are referred to as the *assignment vectors*.

We define the initial tuple of input b it vectors t o b e the rows of 𝒜, *B* = (𝒜^1^, …, *𝒜*^*n*^), where 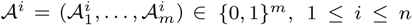. The algorithm starts by constructing the *root* node (*α*_1_, *β*_1_) from the initial set of input vectors *B*_1_ = *B*.

On the *j*^th^ iteration, for a list of input bit vectors

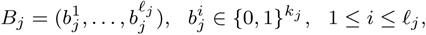

we compute (*α*_*j*_, *β*_*j*_) as follows. First, we compute the longest common prefix *α*_*j*_ := **LCP**(*B*_*j*_) for the bit vectors in *B*_*j*_. Formally, this function is defined as follows,

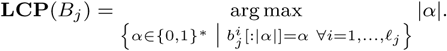

If the computed *α*_*j*_ matches all the input bit vectors, let the assignment vector consist of |*B*_*j*_| zeros, *β*_*j*_ := (0, …, 0) and (*α*_*j*_, *β*_*j*_) is referred to as a *leaf*, which terminates the recursion branch. Otherwise, the assignment vector is set to be the concatenation of next bits in each of the 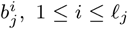 after removing the common prefix *α*_*j*_,

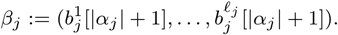

We continue the recursion on the child nodes (*α*_2*j*_, *β*_2*j*_) and (*α*_2*j*+1_, *β*_2*j*+1_), with the new tuples of bit vectors *B*_2*j*_ and *B*_2*j*+1_, respectively, which are defined by partitioning *B*_*j*_ based on the assignments *β*_*j*_ and removing the first |*α*_*j*_| + 1 bits,

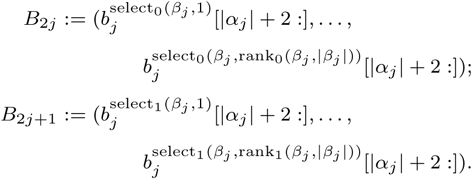

##### Parallel construction via trie merging

To allow for parallel construction, we develop an algorithm to merge wavelet tries constructed on batches of edge colorings that generalizes the methods presented by Grossi and Ottaviano (2012) and Böttcher *et al*. (2017). Merging proceeds by performing an *align* and a *merge* step on each node, starting from the root (Suppl. Figure S-2). Given two wavelet tries *T* ′ and *T* ″ with node sets 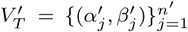 and 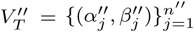 that we want to merge into a new trie *T*, the merging process can be summarized in three steps:

1. **Align:** For the nodes 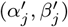 and 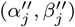, compute the longest common prefix 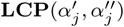, create new nodes with this value and appropriate *β* vectors, and set this to be the parent of the current nodes.
2. **Merge:** Once 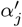 and 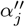 are equal, concatenate 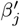 and 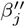.
3. **Repeat:** Move down to *j*’s children and apply the same function until all leaves are reached.

##### Time complexity

Let 𝒜 ∈ {0, 1}^*n*×*m*^ be the annotation matrix. The height of a constructed wavelet trie with nodes *V*_*T*_ depends on the degree to which the input bit vectors share common prefices. Since there can be at most *n* leaves, and the maximum height of the trie is at most *m*, the number of nodes can be at most |*V*_*T*_ | ≤ min(2*n* − 1, 2^*m*^ − 1).

Given two wavelet tries with sets of nodes 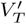 and 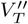, merging is performed in 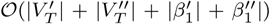 time (Grossi and Ottaviano, 2012). Once a wavelet trie is constructed, queries can be performed in 𝒪 (*h*) time, where *h* ≤ *m* is the height of the trie. To achieve this value, the *β*_*j*_ are compressed with RRR coding (Raman *et al*., 2007) to support rank operations in 𝒪 (1) time.

##### Using prior knowledge to improve compression

One of the most important factors determining compression ratio (see Section 2.4 for a formal definition) of a wavelet trie is the distribution of longest common prefices encountered during construction. We explore whether prior knowledge can be used to form groups of similarly colored edges and help optimize compression ratios.

Given a similarity metric defined on edge colorings, paths can be grouped into classes defined by high similarity between their constituent edge colorings. Then, then class membership can be used to group edges for improved compression. Example class definitions can be based on phylogenetic information (e.g., shared taxonomic IDs) or sequence alignment information (e.g., alignment to a given reference genome).

To encode the assignment of edges to classes, we introduce additional colors and add corresponding new columns (called *class indicator bits*) to the annotation matrix. Additionally, we hypothesize that if the indicator columns are of low index, then edges from the same class are more likely to be co-assigned to matching nodes in a wavelet trie. This would facilitate a partitioning of the rows that has the potential to significantly improve the compression ratio of the wavelet trie. We implement this procedure by providing class information as additional metadata strings, which are then used to augment the coloring of each edge with the color of its corresponding class.

#### 2.3.2 Probabilistic column compression with Bloom filters

For cases where a lossy compression scheme with moderate loss of accuracy will suffice in place of fully lossless compression, we explore a probabilistic compression of the annotation matrix as a near-exact compromise. Since, by definition, the columns of the annotation matrix encode set membership, it is possible to compress them using Bloom filters (Bloom, 1970), a probabilistic data structure for approximate set membership queries.

A *Bloom filter* is a tuple *BF* = (*B, ℋ*), where *B* ∈ {0, 1}^*b*^ is a bit vector and ℋ = {*h*_1_, …, *h*_*d*_} is a collection of *d* hash functions mapping each input to an element of {1, …, *b*}. For simplicity of notation, let **e**_*i*_ ∈ {0, 1}^*b*^ denote a bit vector in which only the *i*^th^ bit is set to one.

Two of the operations supported on this structure are insert and the relation of approximate membership ∈,

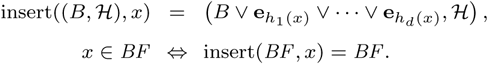

##### Bloom filter reparametrization

Although the Bloom filter has no false negative errors, the *false positive probability* (FPP) of the approximate membership query on a Bloom filter with *s* inserted elements can be approximated (Mitzenmacher, 2001) as

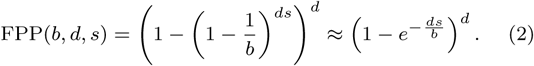

As a corollary, an alternate parametrization of Bloom filters can be derived. Given a target false positive probability *p* and *s* elements to insert, optimal values for *d* and *b* (Mitzenmacher, 2001) *are*

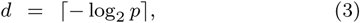

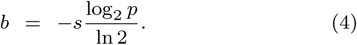

Given an encoding of an annotation matrix 𝒜 ∈ {0, 1}^*n*×*m*^ as a collection of Bloom filters *BF* _1_, …, *BF*_*m*_, the *raw annotation* of an edge *e*_*i*_ ∈ *E* being queried is as follows:

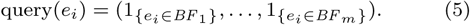

##### Neighborhood-based Bloom filter correction

Following the same rationale as for the wavelet tries, and building on the assumption that edges neighboring in the graph often share a large part of their annotation, we can also drastically improve the compression power of the Bloom filters.

More precisely, given a linear path, we compute the intersection of the colorings of, *e* edges in some neighborhood within the path and obtain an annotation with drastically reduced FPP. If we let 𝒩 (*e*) ⊂ *E* denote the neighborhood of an edge *e* ∈ *E* within a linear path in which all nodes are assumed to share the same annotation, we can then define the *corrected annotation* as

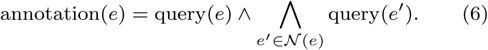

Following the argument in (Mitzenmacher, 2001) (see Formula 2), the FPP for one annotation color of a segment of length, *l* can be approximated as

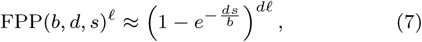

since *l*, false positive errors have to be made to lead the Bloom filter to a false positive error.

This correction method relies on direct access to the underlying graph structure to reference during decoding, in contrast to the wavelet trie approach in which this is not required.

### 2.4 Data

We use several standard datasets to evaluate the performance of our compression schemes. They originate either from viruses (*Virus100, Virus1000*, and *Virus50000*), bacteria (*Lactobacillus*) or humans (*chr22+gnomAD* and *hg19+gnomAD*) and are chosen to test the methods on different coloring distributions, sizes and densities. They further reflect varying graph topologies and allow us to study the effect of topology-informed compression in a robust testbed. We construct de Bruijn graphs of order *k* = 63 for each dataset. We compare the compression performance of the methods by measuring the *compression ratio* for each dataset across methods, defined as the ratio of the number of bits in an annotation matrix and the number of bits in its respective compressed representation. Please refer to Suppl. Section B for a more detailed description of the datasets.

The virus and bacteria datasets are both generated from publicly available GenBank (Clark *et al*., 2016) complete genome data. The *Virus1000* dataset consists of 1000 randomly selected complete virus genomes, whose resulting graph consists of several disjoint, linear paths. The *Virus100* dataset is a subset of this consisting of the first 100 genomes in *Virus1000*. The *Virus50000* dataset consists of the set of 53,412 virus strains present in GenBank on September 30th, 2016, with a greater average similarity between sequences. On these datasets, the colors are defined to be indicators for each genome ID, while the class indicator bits are defined by each genome’s taxonomic genera. The rows of these annotation matrices are very sparse and present degenerate cases with respect to compressability by wavelet tries. As intermediate data points, *Virus3000* and *Virus20000* datasets were constructed as supersets of *Virus1000* (see Suppl. Section B.3 for more details). For brevity, numerical data from these sets is included only in the Supplementary Materials.

The *Lactobacillus* dataset consists of 135 different bacterial strains from the genus *Lactobacillus*, respectively, which leads to a linear topology in the graphs with many shorter paths (*bubbles*) diverging from and reconnecting to the main reference genome paths. The colors are defined to be indicators for genome IDs, while the class indicator bits are defined by each genome’s species.

For the human datasets, the *hg19* assembly of the human reference genome is used as the main reference backbone, together with exome variants from the gnomAD dataset (Consortium *et al*., 2016). The *chr22+gnomAD* dataset is chromosome 22 from this data, resulting in a graph that has a similar structure as the *Virus1000* graph, scaling the number of nodes three-fold and reducing the total number of colors. The *hg19+gnomAD* dataset consists of the human autosomal portion of this data, with a similar topology. On these datasets, the colors are defined to be indicators for the reference chromosomes and the ethnic groups present in the gnomAD data. The colors corresponding to reference chromosomes are designated as the class indicators without adding additional columns (i.e., the sequence variant edges are additionally colored by their corresponding reference chromosome colors).

Table 1 summarizes these collections in terms of their number of nodes and edges for the constructed de Bruijn graphs, as well as their respective numbers of colors and unique colorings derived from the corresponding metadata.

**Table 1.** Datasets used for evaluation. Columns represent number of nodes and edges per dataset, total number of colors and annotations (number of unique edge colorings, or color combinations), and density of the annotation matrices, where the quantity *s* refers to the number of set bits in the annotation matrices.

| Data set | Nodes | Edges $n$ | Colors $m$ | Annotations | Density (%) $\frac{s}{nm}$ |
| --- | --- | --- | --- | --- | --- |
| *Virus100* | 2,954,719 | 2,956,113 | 100 | 463 | 1.056 |
| *Virus1000* | 30,310,634 | 30,347,373 | 1,000 | 11,612 | 0.117 |
| *Virus50000* | 622,587,315 | 625,110,390 | 53,412 | 1,359,843 | 0.006 |
| *Lactobacillus* | 134,951,429 | 135,369,397 | 135 | 6,630 | 1.475 |
| *chr22+gnomAD* | 178,196,890 | 180,023,641 | 9 | 510 | 2.147 |
| *hg19+gnomAD* | 5,714,136,751 | 5,728,489,633 | 30 | 380,051 | 1.762 |

## 3 Evaluation and Applications

In this section, we explore our hypothesis that graph topology can aid in improving compression ratios and study the space complexities of our compression techniques on a variety of viral datasets increasing in size. Finally, we compare the compression ratios of our methods to those of general compression algorithms and those of methods developed specifically for de Bruijn graph color compression.

Experiments were performed on a single thread for Bloom filter compression and ten threads for wavelet trie compression, on the Intel(R) Xeon(R) CPU E5-2697 v4 (2.30GHz) cores of ETH’s shared high-performance compute systems. Run times and peak RAM consumption are reported in Suppl. Table S-7a.

### 3.1 Graph topology affects compression ratios

For both the wavelet trie and Bloom filter compression schemes, we explored methods for encoding graph topology with the goal of improving compression ratios. To this end, we explore the introduction of additional class indicator colors/bits for wavelet tries and graph neighborhood-based annotation correction for Bloom filters.

#### 3.1.1 Class indicator bits significantly improve compression ratio

We test the hypothesis that optimal compression can be achieved by setting class indicator bits in low-index positions in annotation matrices^1^ via an exact test by permuting the annotation matrix column order on the *Virus100* and *Lactobacillus* datasets. More precisely, we generate 100 samples by randomly permuting the columns in the annotation matrix and compress the resulting data to approximate the null distribution of compression file sizes across permutations of the matrix column order (see Figure 2a and Suppl. Figure S-3).

**Fig. 2:**
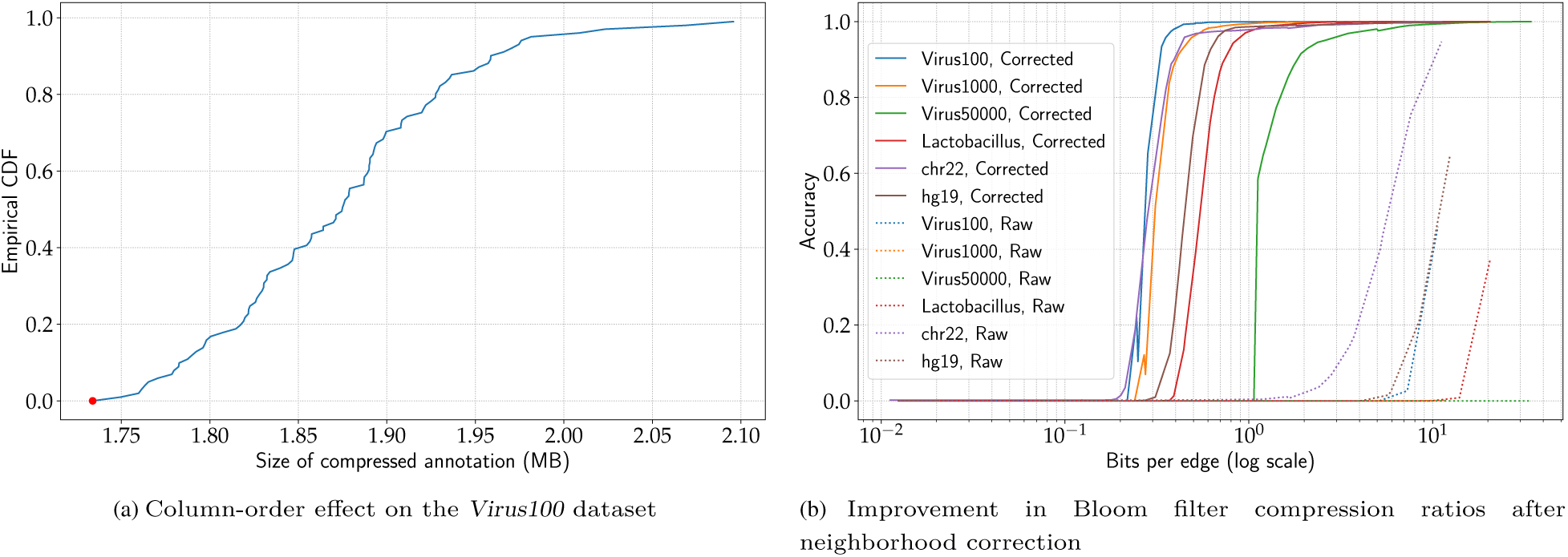
Graph topology improves compression ratios. (a) Distribution of the file sizes of wavelet tries over 100 random permutations of the annotation matrix column order. The original ordering of the columns leads to an optimal file size when class indicator bits are set, as indicated in the CDF by the red dot. (b) Bloom filter decompression accuracy (fraction of correct edge colors) as a function of filter size (bits per edge). Parameters required to achieve 99% accuracy on the uncorrected Bloom filters were not computed.

First, when we test the hypothesis without setting class indicator bits, the compressed file size corresponding to the column ordering induced by the graph construction algorithm is found to not be optimal with respect to its approximated null distribution (see Suppl. Figure S-3). However, when class indicator bits are set in low index positions, the original ordering of columns is optimal with respect to its approximated null distribution, resulting in an empirical *p*-value of *p <* 0.01 (see Figure 2a).

#### 3.1.2 Neighborhood correction improves Bloom filter compression ratio 30- to 70-fold

We study the effects of neighborhood-based Bloom filter correction on all datasets by varying the average number of bits per edge of the Bloom filters and measuring the accuracy of color reconstruction (see Methods, Section 2.3.2). The results show 70-fold decreases in the number of bits required per edge to achieve similar decompression accuracies on almost all datasets (see Figure 2). A notable exception is the *chr22* dataset, where only a 30-fold improvement is observed.

The average number of graph traversal steps needed to correct the Bloom filters to an accuracy of 95% ranges from 99.1 to 207.3 (see Suppl. Table S-1). To correct the Bloom filters to an accuracy of 99%, the average number of traversal steps required ranges from 82.3 to 156.3.

### 3.2 Compression power grows with the number of colors

To test the scalability of the compression methods, we generate a *chain* (a linear hierarchy) of virus graphs ranging from 100 to 1000 randomly selectely selectedd genomes in steps of 100 (i.e., *G*_1_ ⊂ … ⊂ *G*_10_) and measure the compression ratios of the annotations for each graph. On our datasets, the wavelet trie method with class indicator bits set and the Bloom filter method with *FPP <* 0.05 display linear growth in the compression ratio as number of genomes increases to 1000 genomes (see Suppl. Figure S-4), with sublinear growth for more genomes (see Figure 3). Sublinear growth is observed in the wavelet trie method without class indicator bits and, to a lesser extent, the Bloom filter method with *FPP <* 0.01 (see Figure 3 and Suppl. Figure S-4). A two-fold decrease in compression ratio is observed when the false positive probability criterion for the Bloom filters is decreased from 0.05 to 0.01.

**Fig. 3:**
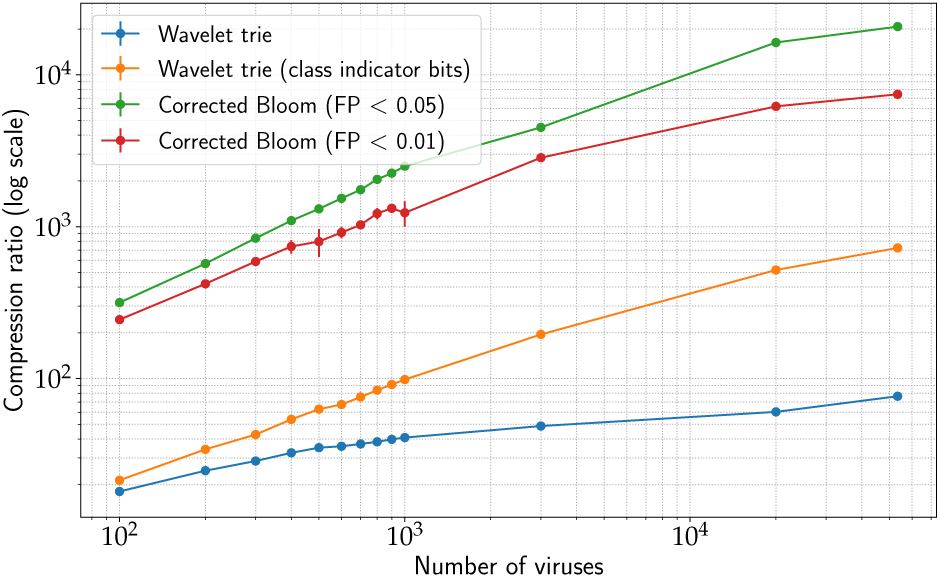
Growth of compression ratios. Compression ratios on virus graphs of increasing genome count. Error bars were computed from the virus graph chains resulting from six random draws of the *Virus1000* dataset (see Section 3.2.

### 3.3 Wavelet tries and Bloom filters improve on state-of-the-art compression ratios

Finally, we close with a side-by-side comparison of the various de Bruijn graph color compression schemes presented in Section 1. In addition to these domain-specific methods, we include two popular general-purpose static compression methods, *gzip* and *bzip2. gzip* is an implementation of the *LZ77* algorithm that encodes blocks of text, while *bzip2* performs a sequence of transformations, including run-length encoding, BWT, move-to-front transforms, and Huffman coding.

Table 2 lists the number of bits required per edge to compress our experimental collections.

**Table 2.** Compression performance of wavelet trie and Bloom filter schemes (measured as number of bits per edge). Each dataset is encoded with eight different compression schemes, including general compression algorithms such as *gzip* and *bzip2*, existing methods specific to colored de Bruijn graphs such as *VARI* (Muggli et al., 2017) *and Rainbowfish (RBF, Almodaresi* et al. *(2017)), as well as the wavelet trie encoding with (WTr (CI)*) and without (*WTr*) the class indicator bits set, and the Bloom filter compression at *>* 95% (*BF 95%*) and *>* 99% (*BF 99%*) accuracy. All compression rates are measured as average number of bits per edge. VARI numbers correspond to compilation with 1024 bit support. *: On these datasets, VARI and RBF results are generated by exporting the annotation data in compatible formats. ^*†*^ : class indicator bits set in all figures. ‡: Consumed more than 400GB memory limit.

| Data set | Colors<br>$m$ | gzip | bzip2 | VARI | RBF | Proposed | | | |
| --- | --- | --- | --- | --- | --- | --- | --- | --- | --- |
|  |  |  |  |  |  | WTr | WTr<br>(CI) | BF 95% | BF<br>99.0% |
| *Virus100* | 100 | 11.4 | 4.8 | 9.8 | 5.8 | 5.7 | 4.9 | 0.36 | 0.44 |
| *Virus1000* | 1,000 | 26.5 | 7.5 | 14.7 | 9.7 | 23.8 | 10.5 | 0.49 | 0.82 |
| *Virus50000* | 53,412 | 135.3 | 37.7 | 56.0* | ‡* | 698.4 | 73.7 | 2.58 | 7.41 |
| *Lactobacillus* | 135 | 15.6 | 5.7 | 19.3 | 7.8 | 7.3 | 5.6 | 0.95 | 1.40 |
| *chr22+gnomAD*<sup>†</sup> | 9 | 4.6 | 2.7 | 17.3* | 3.3* | N/A | 2.4 | 0.45 | 2.41 |
| *hg19+gnomAD*<sup>†</sup> | 30 | 10.9 | 5.4 | 14.5* | 5.6* | N/A | 5.4 | 0.68 | 1.82 |

#### 3.3.1 Wavelet trie compression ratios match state-of-the-art

Our results show that wavelet trie compression outperforms *gzip* and the *VARI* method on most datasets, while performing marginally better than *Rainbowfish* and marginally worse than *bzip2* (see Table 2). The *Virus100, Virus1000, Virus50000*, and *Lactobacillus* datasets are compressed to 5.8, 23.8, 698.4, and 7.3 bits per edge, respectively. The Virus1000 and Virus50000 datasets are notable in that wavelet tries without indicator bits set exhibit the worst compression performance among the tested methods. Setting class indicator bits leads to a two-fold improvement in the compression performance on the Virus1000 dataset (from 23.8 bits per edge to 10.5), ten-fold improvement on the *Virus50000* dataset (from 698.4 to 73.7 bits per edge), and marginal improvements in performance on the other datasets (4.9 and 5.6 bits per edge on the *Virus100* and *Lactobacillus* datasets, respectively). In this setting, the *chr22+gnomAD* and *hg19+gnomAD* datasets are compressed to 2.4 and 5.5 bits per edge.

#### 3.3.2 Bloom filters improve on state-of-the-art by an order of magnitude

At an accuracy of 95%, our method is considerably more space efficient, achieving compression ratios over an order of magnitude greater than *bzip2* and *Rainbowfish*. An average of 0.35 and 0.49 bits per edge are required to compress the *Virus100* and *Virus1000* datasets, respectively, compared to 5.8 and 9.7 bits for *Rainbowfish* and 4.8 and 7.5 bits for *bzip2*. An average of 2.4 bits per edge are required to compress the *Virus50000* data set, compared to 37.7 bits for *bzip2*. We were unable to compress this dataset using the *Rainbowfish* method due to its RAM consumption exceeding the per-job limit on our computing system. On the *Lactobacillus* dataset, an average of 1 bit per edge are required, compared to 7.8 bits for *Rainbowfish* and 5.7 bits for *bzip2*. On the *chr22+gnomAD* and *hg19+gnomAD* datasets, 0.45 and 0.68 bits are required per edge, compared to 2.7 and 5.4 bits for *bzip2*, and 3.3 and 5.6 bits for *Rainbowfish*.

At 99% accuracy, an increasing number of bits are required per edge with increased virus dataset size (see Table 2). Fold-increases in the number of bits per edge from 1.3 bits (*Virus100*) to 5.4 bits (*chr22+gnomAD*) are required.

## 4 Discussion

In this study, we have addressed the problem of encoding metadata as edge colors of a given graph and demonstrated its application to de Bruijn graphs by presenting two distinct compression schemes. First, we have developed a novel application and extended parallel construction method of the wavelet trie data structure on general sequences of bit vectors that employs an iterative merging scheme to build larger tries from many smaller instances. Further, we have presented a probabilistic, compressed representation using approximate set representations that can store an arbitrary amount of annotations on the graph and allows for greater compression ratios by taking advantage of information shared between neighboring nodes to correct errors.

We have shown that utilizing the topology of the underlying graph helps in achieving improved compression rates. For the wavelet tries, we used indicators for the backbone regions of the de Bruijn graph positioned in low-index columns of the annotation matrix and for the Bloom filter approach, we used neighboring linear regions in the graph for error correction.

Either representation can be efficiently decompressed and queried to retrieve the coloring of arbitrary paths in the graph. Although it is helpful to know the frequency of individual colors upfront to design an optimal order of columns for the wavelet trie compression or to optimally choose the size of the individual Bloom filters used, these parameters can be easily estimated from a subsample of the input data, allowing to directly build the full coloring.

We have shown the utility of our approaches on different biological datasets, including data from virus, bacteria and human genomes, representing different classes of graph topologies and colorings. On all datasets we achieve comparable or strongly increased compression performance at very high levels of decompression accuracy. Notably, our approach is dynamic and allows for an easy extension with additional labels/colors or for changes in the underlying graph structures, enabling the augmentation of large colored graphs with new annotations — a scenario commonly occurring in the genomics setting. Additionally, the wavelet trie model is fully dynamic, allowing for label and edge removal.

A possible limitation of the wavelet trie method is its reliance on shared segments (contiguous subsequences), especially in the first few columns of the annotation matrix, to effectively partition the rows for optimal compression. The results on the viral datasets confirm that, given an annotation matrix with very sparse and mutually-exclusive rows, wavelet tries underperform relative to other methods due to tree imbalance. While this is partially addressed by setting class indicator bits in the annotation matrix, a more principled approach with less user input will become necessary in future work and could involve an analysis of the de Bruijn graph topology to algorithmically determine optimal backbone paths. Further improvements in compression ratio could be gained by an optimal ordering of the rows of the annotation matrix, but at the additional cost of maintaining a map from graph coordinates to their respective annotation matrix rows.

One of the limitations of our Bloom filter correction method is its reliance on the presence of long, identically-colored paths for correction. While this assumption worked well for the *Virus100* and *Virus1000* datasets, the shorter linear paths in the larger sets reduced our ability to correct errors in this fashion. Despite its higher compression ratio, one restriction of the Bloom filter-based method is that its corresponding graph must be accessible for reference. Although this is already done in our application, it couples annotation query times to graph query times. To decouple the graph from the filters, an additional structure could be constructed to indicate edges in the graph at which changes in coloring occur. Such a structure would then allow for the assumption that colors remain constant in linear regions to be relaxed.

Future work on probabilistic compression will focus on improving scaling properties. In a dynamic setting, if a dataset grows rapidly in the number of edges, the decoding accuracy will eventually drop, ultimately requiring a re-initialization into a larger Bloom filter. Further, despite being dynamic, the current probabilistic representation does not allow for the removal edges from the graph. To support this, we could replace the Bloom filters with other probabilistic set representations that allow for item removal (Bender *et al*., 2012; Fan *et al*., 2014). Lastly, an additional space improvement could be achieved with more space efficient probabilistic set representations such as compressed Bloom filters (Mitzenmacher, 2001).

## Acknowledgments

The authors would like to thank Torsten Hoefler as well as all members of the Biomedical Informatics group at ETH Zurich, in particular Amir Joudaki, Viktor Gal, and Gideon Dresdner, for valuable discussions, questions and feedback. Carsten Eickhoff is funded by the Swiss National Science Foundation Ambizione Program under grant agreement no. 174025. Harun Mustafa and Mikhail Karasikov are funded by the Swiss National Science Foundation grant #407540_167331 “Scalable Genome Graph Data Structures for Metagenomics and Genome Annotation” as part of Swiss National Research Programme (NRP) 75 “Big Data”.

*The authors declare no conflicts of interest*.

## Footnotes

1 *H*_0_: column ordering does not influence compression ratios when class indicator bits are set

