## Supplementary Materials for "Dynamic compression schemes for graph coloring"

[1]Department of Computer Science, ETH Zurich, Zurich, Switzerland
[2]University Hospital Zurich, Biomedical Informatics Research, Zurich, Switzerland
[3]SIB Swiss Institute of Bioinformatics, Zurich, Switzerland
[4]Brown Center for Biomedical Informatics, Brown University, Providence, RI, USA
[‡]Equal contribution. [*]To whom correspondence should be addressed.

### A Parallel wavelet trie construction

For this method, we define the *descendants* function $\mathbf{D} : \{1,\ldots,|V|\} \to 2^{\{1,\ldots,|V|\}}$ for the wavelet trie $T = (V, E)$ with nodes $V = \{(\alpha_j, \beta_j)\}_{j=1}^{n}$ by the recurrence

$$j \in \mathbf{D}(j),\ \forall j \in \{1,\ldots,n\},$$
$$k \in \mathbf{D}(j) \text{ and } \beta_k \neq \varepsilon \Rightarrow \{2k, 2k+1\} \subset \mathbf{D}(j).$$

The three steps in the merging operations are as follows:

#### A.1 Align

Given nodes $(\alpha'_j, \beta'_j)$ and $(\alpha''_j, \beta''_j)$, we compute their longest common prefix

$$\alpha_j \leftarrow \mathbf{LCP}(\{\alpha'_j, \alpha''_j\}).$$

If $\alpha_j \neq \alpha'_j$, we let

$$\hat{\beta}'_j \leftarrow \underbrace{\alpha'_j[|\alpha_j|+1]\cdots\alpha'_j[|\alpha_j|+1]}_{|\beta'_j|}$$

and update the indices in $T'$ by applying the transformation $j \leftarrow 2j + \alpha'_j[|\alpha_j|+1]$ and updating all nodes $k \in \mathbf{D}(j)$ accordingly. We then let $\alpha'_j \leftarrow \alpha_j$ and $\beta'_j \leftarrow \hat{\beta}'_j$ and truncate the prefix in the newly created child nodes,

$$\alpha'_{2j} \leftarrow \alpha'_{2j}[|\alpha_j|+2:] \qquad \text{if } \hat{\beta}'_j[1] = 0,$$
$$\alpha'_{2j+1} \leftarrow \alpha'_{2j+1}[|\alpha_j|+2:] \qquad \text{if } \hat{\beta}'_j[1] = 1.$$

If $\alpha_j \neq \alpha''_j$, the second trie is processed accordingly.

#### A.2 Merge

If $\mathrm{rank}_1(\beta'_j, |\beta'_j|) = 0$ and $\mathrm{rank}_1(\beta''_j, |\beta''_j|) = 0$, then terminate. Otherwise, merge the two assignment vectors

$$\beta_j \leftarrow \beta'_j \beta''_j$$

#### A.3 Repeat

The merging algorithm is then performed on nodes $n_{2j}$ and $n_{2j+1}$ depth-first to continue the recursion.

If two wavelet tries constructed from bit vectors of different lengths are merged, this merging algorithm leads to the decoding of bit vectors with trailing zeros. Since we intend to use these vectors as indicators for various metadata, the presence of extra trailing zeros in the decoded bit vector does not represent false information.

### B Data used for evaluation

#### B.1 *Lactobacillus*

This dataset is composed of 135 strains of bacteria in GenBank Clark *et al.* (2016) from the *Lactobacillus* genus. The columns in this graph's annotation matrix indicate presence of an edge in each of the strains. Because of the low variability in the input sequences, they are represented as a graph with a predominantly linear topology and short variant paths (called *bubbles*). One genome from each of the species was chosen as a backbone path. The resulting graph had 134,951,429 unique $k$-mers, 135,369,397 edges, and 6,630 unique edge colorings (i.e., bit combinations). See Supplementary Section C for a list of the bacterial strains used.

#### B.2 Virus50000

This dataset consists of 53,412 viral genome sequences downloaded from GenBank via the eUtils API on 09/30/2016. The search term applied was `txid10239 [orgn] AND "complete genome" [Title] NOT txid131567 [orgn]`. The data represents a set of sequences with both high variability to each other but also good pairwise conservation for subspecies. It reflects a wide range of variability and provides a good testing bed for the application within a colored de Bruijn graph setting. The graph corresponding to this dataset contains 622,587,315 nodes, 625,110,390 edges, and 1,359,843 unique edge colorings.

#### B.3 Virus1000, Virus3000, Virus20000

This dataset is composed of 1000 virus genomes randomly selected from the *Virus50000* set, meant to study a graph whose topology is a series of almost mutually-exclusive loops with slight variation. The columns in this graph's annotation matrix indicate presence of edges in each of the virus genomes. Similar to the *Lactobacillus* dataset, the viruses were grouped by the first word of their names and the first species in each group was assigned as a backbone path. The resulting annotation bit matrix is very sparse and adjacent rows are either almost identical or almost mutually exclusive. This graph contains 30,310,634 unique $k$-mers, 30,347,373 edges, and 11,612 unique edge colorings.

The *Virus3000* and *Virus20000* datasets are supersets of *Virus1000*. The *Virus3000* graph contains 82,418,835 unique $k$-mers, 82,579,519 edges, and 52,187 unique edge colorings. The *Virus20000* graph contains 357,552,076 unique $k$-mers, 358,683,520 edges, and 537,344 unique edge colorings.

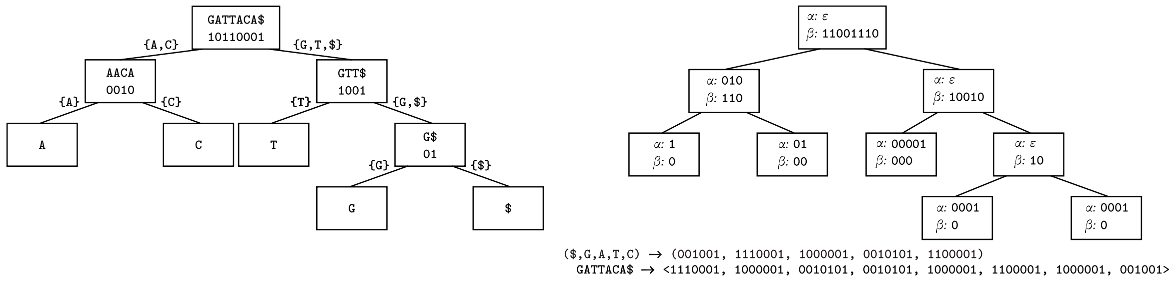


Fig. S-1: **Comparison of a wavelet tree (left) and a wavelet trie (right) constructed for the string `GATTACA$` and the binary encodings of its characters, respectively. Wavelet tree:** Characters for each node of the wavelet trie are divided equally amongst its two children, with a bit vector indicating these assignments. A node becomes a leaf when it is assigned only one character. In internal nodes, only the bit vectors need to be stored to allow traversal to the correct leaf when querying the wavelet tree. **Wavelet trie:** In a wavelet trie, strings are encoded as tuples of bit vectors. At a node, the common prefix of the bit vectors is extracted and the next significant bit is used to assign the bit vector suffices to that node's children. A node becomes a leaf when all bit vectors assigned to it are equal. In both structures, index queries are resolved by traversing the tree and performing rank operations on the assignment bit vectors. In this example, ASCII codes are used to define the binary codes for each character.

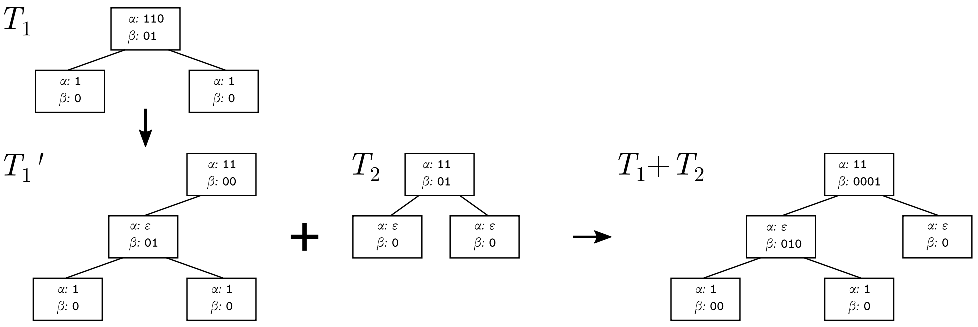


Fig. S-2: **Merging of wavelet tries $T_1$ and $T_2$ to form the wavelet trie $T_1 + T_2$.** Starting from the root node, the common prefix of the two $\alpha$ vectors is found and new $\beta$ vectors are computed from the their remainders. These become new parent nodes and the initial nodes' $\alpha$ vectors are updated to their respective remainders after removing the common prefix (e.g., the conversion from from $T_1$ to $T_1'$). When the two $\alpha$s are equal, their respective $\beta$s are concatenated and the merging function is applied to their children. When a leaf is reached in one tree, but the equivalent node in the other tree is internal (e.g., the left child of the root in $T_1 + T_2$), the leaf is merged by appending or prepending additional zeros to the $\beta$ vectors of all left ancestors. Note that extra leaf nodes producing trailing zeros in the decoded bit vectors are added during the merging process. See Section 2.3.1 for more details.

#### B.4 Virus100

This is a subset of the Virus1000 set containing only 100 virus strains used to facilitate the permutation tests in Section 3.1.1. This graph contains 2,954,719 unique $k$-mers, 2,956,113 edges, and 463 unique edge colorings.

#### B.5 chr22+gnomAD

This graph consists of chromosome 22 from the `hg19` assembly of the human reference genome as the main reference backbone. To provide genetic variability, the set of exome variants from the gnomAD dataset were incorporated into the graph Lek *et al.* (2016). This larger dataset is meant to analyze the properties of the trie when the underlying graph is large, but with little variability. The columns in this graph's annotation matrix are defined as indicators for its edges' presence in 9 ethnic groups defined in the dataset. The first column in the matrix is used to indicate edges which are present in the reference genome and serves as the backbone bit. The graph contains 178,196,890 unique $k$-mers, 180,023,641 edges, and 510 unique edge colorings.

#### B.6 hg19+gnomAD

This graph was constructed from the same datasets as the one described above, using data from the full human autosome. The same definition is used for the annotation matrix columns, with 9 columns being used to indicate edges observed in the defined ethnic groups and 22 prefix columns being used to indicate presence in the first 22 reference chromosomes as the backbone bits. This graph's topology was designed to be analogous to the Virus1000 dataset, but with 1000× the number of rows and one-tenth of the number of annotation columns. It contains 5,714,136,751 unique $k$-mers, 5,728,489,633 edges, and 380,051 unique edge colorings.

### C List of bacterial strains used

- *Lactobacillus acidophilus*
  - 30SC chromosome, complete genome
  - 30SC plasmid pRKC30SC2, complete sequence
  - La-14, complete genome
  - NCFM chromosome, complete genome

- *Lactobacillus amylovorus*
  GRL 1112 chromosome, complete genome
  GRL 1112 plasmid1, complete sequence
  GRL 1112 plasmid2, complete sequence
  GRL1118 chromosome, complete genome
  GRL1118 plasmid1, complete sequence
  GRL1118 plasmid2, complete sequence
- *Lactobacillus brevis*
  ATCC 367, complete genome
  ATCC 367 plasmid 1, complete sequence
  ATCC 367 plasmid 2, complete sequence
  KB290 DNA, complete genome
  KB290 plasmid pKB290-1 DNA, complete genome
  KB290 plasmid pKB290-2 DNA, complete genome
  KB290 plasmid pKB290-4 DNA, complete genome
  KB290 plasmid pKB290-5 DNA, complete genome
  KB290 plasmid pKB290-7 DNA, complete genome
  KB290 plasmid pKB290-9 DNA, complete genome
  KB290 plasmid pKB290-3 DNA, complete genome
  KB290 plasmid pKB290-6 DNA, complete genome
  KB290 plasmid pKB290-8 DNA, complete genome
- *Lactobacillus buchneri*
  NRRL B-30929 plasmid pLBUC01, complete sequence
  NRRL B-30929 plasmid pLBUC03, complete sequence
  NRRL B-30929 chromosome, complete genome
  NRRL B-30929 plasmid pLBUC02, complete sequence
  CD034 plasmid pCD034-2, complete sequence
  CD034 plasmid pCD034-1, complete sequence
  CD034 chromosome, complete genome
  CD034 plasmid pCD034-3, complete sequence
- *Lactobacillus casei*
  ATCC 334 plasmid 1, complete sequence
  ATCC 334 chromosome, complete genome
  BD-II chromosome, complete genome
  BD-II plasmid pBD-II, complete sequence
  BL23 chromosome, complete genome
  LC2W chromosome, complete genome
  LC2W plasmid pLC2W, complete sequence
  LOCK919, complete genome
  LOCK919 plasmid pLOCK919, complete sequence
  W56, complete genome
  W56 plasmid pW56, complete sequence
  str. Zhang plasmid plca36, complete sequence
  str. Zhang chromosome, complete genome
- *Lactobacillus crispatus* ST1, complete genome
- *Lactobacillus delbrueckii subsp. bulgaricus*
  2038 chromosome, complete genome
  ATCC 11842 chromosome, complete genome
  ATCC BAA-365 chromosome, complete genome
  ND02 chromosome, complete genome
  ND02 plasmid unnamed, complete sequence
- *Lactobacillus fermentum*
  CECT 5716 chromosome, complete genome
  F-6, complete genome
  IFO 3956, complete genome
- *Lactobacillus gasseri* ATCC 33323 chromosome, complete genome
- *Lactobacillus helveticus*
  CNRZ32, complete genome
  DPC 4571, complete genome
  H10 chromosome, complete genome
  H10 plasmid pH10, complete sequence
  R0052 chromosome, complete genome
- *Lactobacillus johnsonii*
  DPC 6026 chromosome, complete genome
  FI9785 plasmid p9785S, complete sequence
  FI9785 chromosome, complete genome
  FI9785 plasmid p9785L, complete sequence
  N6.2, complete genome
  NCC 533, complete genome
- *Lactobacillus kefiranofaciens*
  ZW3 plasmid pWW1, complete sequence
  ZW3 chromosome, complete genome
  ZW3 plasmid pWW2, complete sequence
- *Lactobacillus paracasei subsp. paracasei*
  8700:2, complete genome
  8700:2 plasmid 1, complete sequence
  8700:2 plasmid 2, complete sequence
- *Lactobacillus plantarum*
  16, complete genome
  16 plasmid Lp16A, complete sequence
  16 plasmid Lp16C, complete sequence
  16 plasmid Lp16E, complete sequence
  16 plasmid Lp16F, complete sequence
  16 plasmid Lp16H, complete sequence
  16 plasmid Lp16L, complete sequence
  16 plasmid Lp16B, complete sequence
  16 plasmid Lp16D, complete sequence
  16 plasmid Lp16G, complete sequence
  16 plasmid Lp16I, complete sequence
  JDM1, complete genome
  subsp. plantarum P-8, complete genome
  subsp. plantarum P-8 plasmid LBPp2, complete sequence
  subsp. plantarum P-8 plasmid LBPp3, complete sequence
  subsp. plantarum P-8 plasmid LBPp5, complete sequence
  subsp. plantarum P-8 plasmid LBPp6, complete sequence
  subsp. plantarum P-8 plasmid LBPp1, complete sequence
  subsp. plantarum P-8 plasmid LBPp4, complete sequence
  subsp. plantarum ST-III chromosome, complete genome
  subsp. plantarum ST-III plasmid pST-III, complete sequence
  WCFS1, complete genome
  WCFS1 plasmid pWCFS101, complete sequence
  WCFS1 plasmid pWCFS102, complete sequence
  WCFS1 plasmid pWCFS103, complete sequence
  ZJ316, complete genome
  ZJ316 plasmid pLP-ZJ101, complete sequence
  ZJ316 plasmid pLP-ZJ102, complete sequence
  ZJ316 plasmid pLP-ZJ103, complete sequence
- *Lactobacillus reuteri*
  DSM 20016 chromosome, complete genome
  I5007, complete genome
  I5007 plasmid pLRI03, complete sequence
  I5007 plasmid pLRI02, complete sequence
  I5007 plasmid pLRI05, complete sequence
  I5007 plasmid pLRI06, complete sequence
  I5007 plasmid pLRI01, complete sequence
  I5007 plasmid pLRI04, complete sequence
  JCM 1112, complete genome
  SD2112 chromosome, complete genome
  SD2112 plasmid pLR585, complete sequence
  SD2112 plasmid pLR580, complete sequence
  SD2112 plasmid pLR581, complete sequence
  SD2112 plasmid pLR584, complete sequence

TD1, complete genome

- *Lactobacillus rhamnosus*
  ATCC 8530 chromosome, complete genome
  GG, complete genome
  GG chromosome, complete genome
  Lc 705 chromosome, complete genome
  Lc 705 plasmid pLC1, complete sequence
  LOCK900, complete genome
  LOCK908, complete genome
- *Lactobacillus ruminis* ATCC 27782 chromosome, complete genome
- *Lactobacillus sakei* subsp. sakei 23K chromosome, complete genome
- *Lactobacillus salivarius*
  CECT 5713 plasmid pHN1, complete sequence
  CECT 5713 plasmid pHN2, complete sequence
  CECT 5713 chromosome, complete genome
  CECT 5713 plasmid pHN3, complete sequence
  UCC118 plasmid pSF118-20, complete sequence
  UCC118 plasmid pSF118-44, complete sequence
  UCC118 chromosome, complete genome
  UCC118 plasmid pMP118, complete sequence
- *Lactobacillus sanfranciscensis*
  TMW 1.1304 chromosome, complete genome
  TMW 1.1304 plasmid pLS1, complete sequence
  TMW 1.1304 plasmid pLS2, complete sequence

### D List of viral strains used

See the GitHub repository for a list of viral strains used.

### E Supplementary Figures and Tables

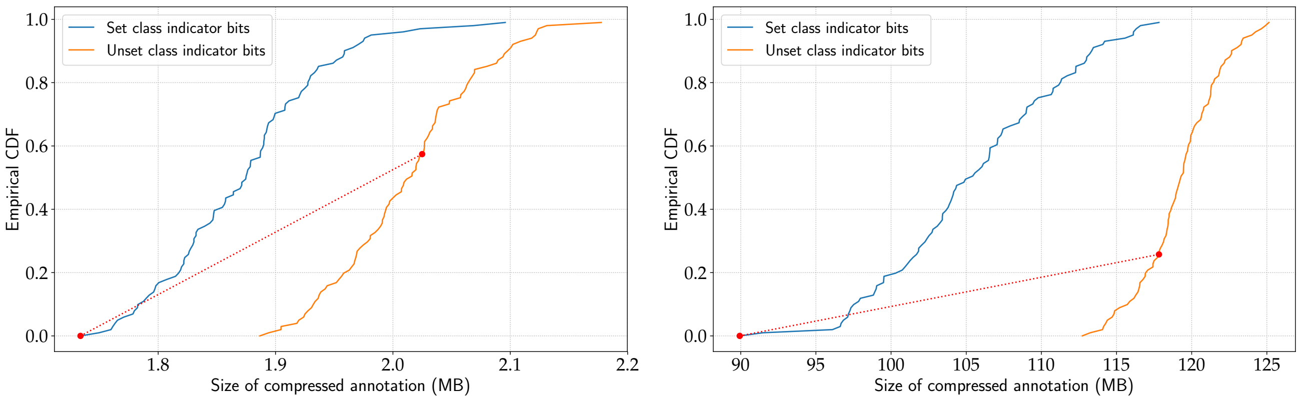


(a) *Virus100* **dataset** (b) *Lactobacillus* **dataset**

Fig. S-3: **Distributions of the file sizes of wavelet tries over 100 random permutations of the** *Virus100* **and** *Lactobacillus* **annotation matrix column orders.** The red lines indicate the mapping between the positions in the CDFs of the original column orderings. Setting class indicator (CI) bits in both datasets leads to decreases in the sizes of the compressed files, with the original ordering being optimal when CI bits are set. The original orderings are not optimal when CI bits are not set. In both datasets, the CDFs are not disjoint, so there exist permutations of the columns with CI bits that exhibit worse compression performance than some permutations without CI bits set.

| | **Bloom filter ($FPP < 0.05$)** | | **Bloom filter ($FPP < 0.01$)** | | **Wavelet trie** | **Wavelet trie CI** |
|---|---|---|---|---|---|---|
| **Data set** | **Query time (ms)** | **Neighborhood** | **Query time (ms)** | **Neighborhood** | **Query time (ms)** | **Query time (ms)** |
| *Virus100* | 1.575 | 207.333 | 1.338 | 156.261 | 0.014 | 0.015 |
| *Virus1000* | 5.571 | 175.155 | 4.795 | 122.594 | 0.114 | 0.055 |
| *Virus50000* | 430.59 | 117.655 | 591.609 | 96.271 | 36.592 | 1.352 |
| *Lactobacillus* | 2.051 | 124.688 | 1.834 | 105.560 | 0.025 | 0.019 |
| *chr22+gnomAD* | 1.085 | 99.121 | 1.406 | 82.272 | N/A | 0.007 |
| *hg19+gnomAD* | 1.972 | 124.618 | 2.168 | 98.259 | N/A | 0.014 |

Table S-1. Average query time(ms) and distance traversed

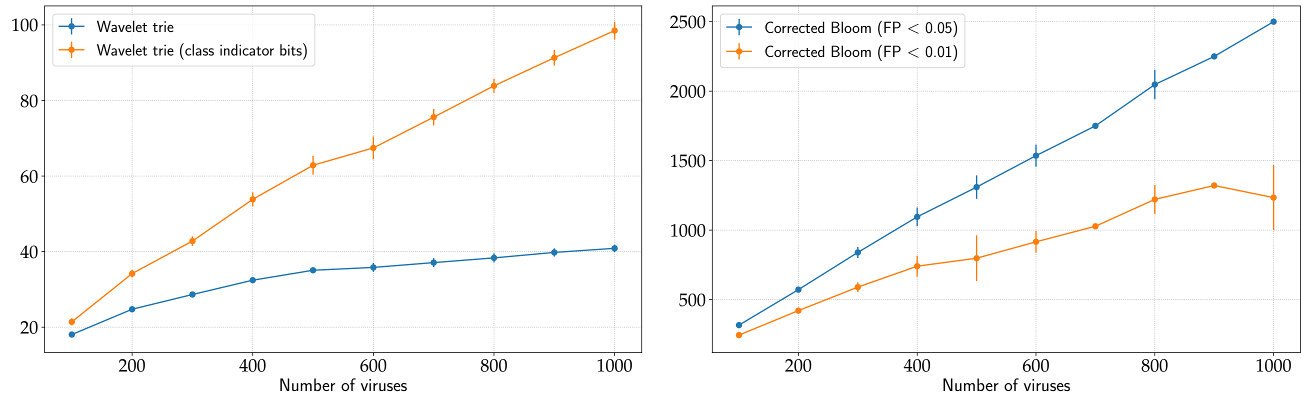


(a) (b)

Fig. S-4: **Growth of compression ratios on the *Virus100* to *Virus1000* datasets.**

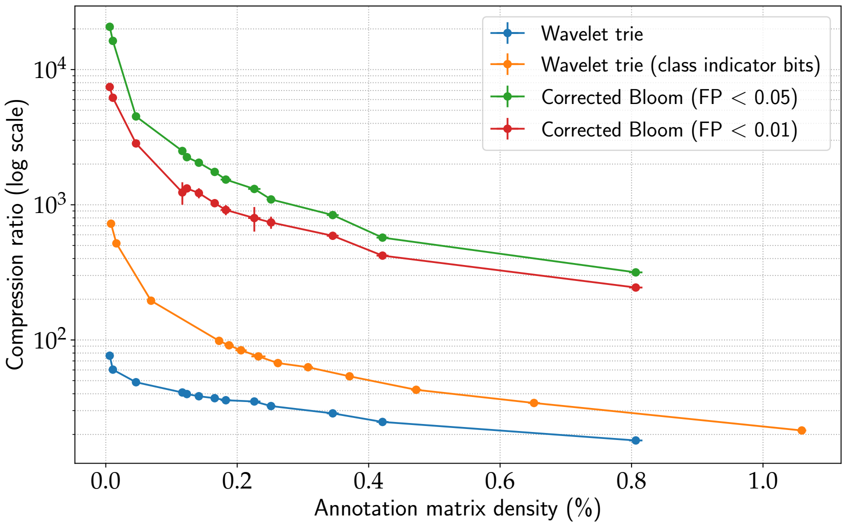


Fig. S-5: **Decrease in compression ratio with increasing annotation matrix density on the virus data sets.** Each data point represents a mean value for across the genomes of a virus collection of a given size. Error bars in both axis represent standard deviation.

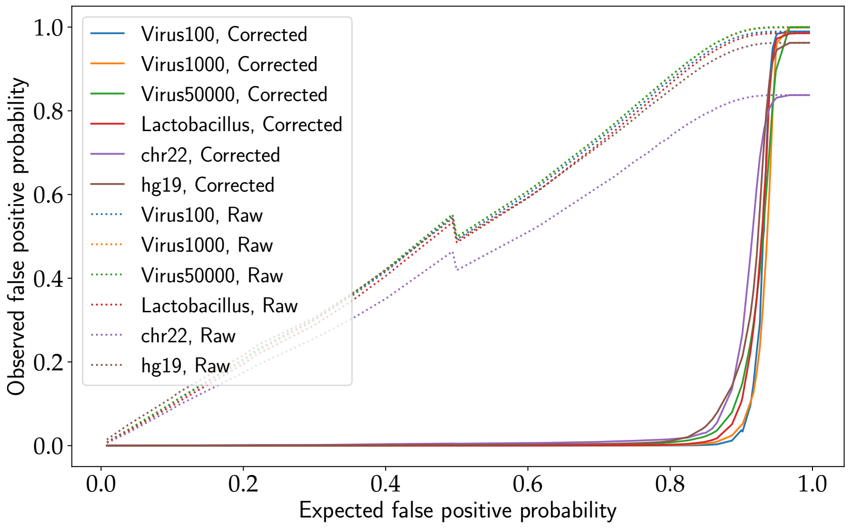


Fig. S-6: **Observed vs. expected per-bit false positive probabilities of the Bloom filters before and after correction.** The drop in the observed false positive probability at $E[FPP] = 0.5$ can be explained by the change in optimal number of hash functions at that point.

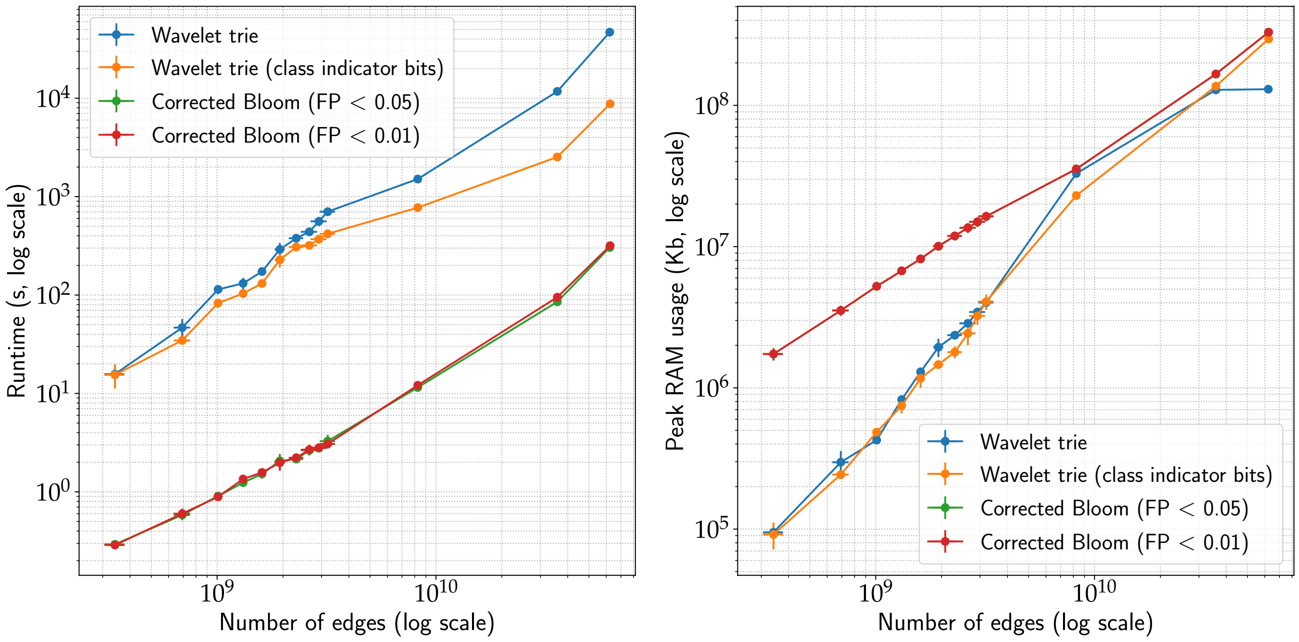


| | **Bloom filter ($FPP < 0.05$)** | | **Bloom filter ($FPP < 0.01$)** | | **Wavelet trie** | | **Wavelet trie CI** | |
|---|---|---|---|---|---|---|---|---|
| **Data set** | **Time (s)** | **RAM (Kb)** | **Time (s)** | **RAM (Kb)** | **Time (s)** | **RAM (Kb)** | **Time (s)** | **RAM (Kb)** |
| *Virus100* | 0.260 | 1,517,176 | 0.242 | 1,517,184 | 6.02 | 84,320 | 5.14 | 69,160 |
| *Virus1000* | 3.029 | 15,523,744 | 2.935 | 15,524,912 | 203.56 | 3,906,892 | 197.44 | 3,793,544 |
| *Virus50000* | 304.352 | 329,415,180 | 317.475 | 329,783,772 | 46,914.00 | 130,183,272 | 8,774.00 | 295,458,816 |
| *Lactobacillus* | 18.557 | 74,243,264 | 19.121 | 74,249,580 | 262.16 | 4,762,584 | 198.52 | 2,732,904 |
| *chr22+gnomAD* | 37.408 | 77,010,668 | 64.773 | 77,032,644 | N/A | N/A | 190.18 | 2,330,828 |
| *hg19+gnomAD** | 1,009.93 | 16,572,464 | 1,431.11 | 17,390,696 | N/A | N/A | 9,769.00 | 149,505,056 |

(a) **Summary of runtimes and maximum memory usage.** Bloom filter runtimes are for a single thread, while wavelet trie runtimes are for ten threads. *: performed with succinct graph representation.

Fig. S-7: **Runtime and peak RAM usage on the virus data sets.** In the plots, each data point represents a mean value across the genomes of each virus collection of a given size (e.g., six draws of the *Virus100* dataset are averaged in both axes and represented by one point). Error bars in both axes represent standard deviation.
